# Temperature-driven biogeography of marine giant viruses infecting the picoeukaryote *Micromonas*

**DOI:** 10.1101/2024.12.12.627685

**Authors:** David Demory, Hisashi Endo, Anne-Claire Baudoux, Estelle Bigeard, Nigel Grimsley, Nathalie Simon, Hiroyuki Ogata, Joshua S. Weitz

**Affiliations:** Sorbonne Université, CNRS, USR 3579 Laboratoire de Biodiversité et Biotechnologies Microbiennes (LBBM), Observatoire Océanologique, Banyuls-sur-Mer, F-66650, France; Bioinformatics Center, Institute for chemical Research, Kyoto University, Uji, Japan; Sorbonne Université, CNRS, UMR7144 Adaptation et Diversité en Milieu Marin (ECOMAP), Station Biologique de Roscoff, F-29680, France; Department of Biology, University of Maryland, College Park, Maryland, USA; Department of Physics, University of Maryland, College Park, Maryland, USA; University of Maryland Institute for Health Computing, North Bethesda, Maryland, USA

**Keywords:** Marine viruses, phytoplankton, temperature, Biogeography, Micromonas, TARA, ecological modeling

## Abstract

Climate shapes the biogeography of microbial and viral communities in the ocean. Among abiotic factors, temperature is one of the main drivers of microbial community distribution. However, we lack knowledge on how temperature shapes the life history traits, population dynamics, and the biogeography of marine viruses. This study integrates mathematical modeling with in situ observations to investigate the temperature-driven biogeography of marine viruses. We focused on prasinoviruses, a group of giant viruses that infect the picoeukaryote *Micromonas*, a widespread phytoplankton with thermotypes adapted from poles to tropics. Analyzing the Tara Oceans and Polar Circle databases, we found that temperature is the primary determinant of *Micromonas* virus (MicV) distribution in the surface ocean. Phylogenetic reconstruction of MicV-s revealed that these viruses form several groups with cryophile or cryo-mesophile preferences. We applied a mechanistic model to describe temperature-driven population dynamics, allowing us to predict the global presence and absence of MicV-s. Lysis and infection emerged as reliable predictors of MicV distribution, indicating that temperature-driven cellular mechanisms significantly shape viral community structure and distribution. This study provides new insights into the role of temperature in regulating viral populations, emphasizing the importance of integrating modeling with observations to understand marine viral ecology.

## 1 Introduction

Viruses are drivers of microbial dynamics and regulate both ecosystems and biogeochemical cycles in the oceans (1; 2). Among marine viruses, the phylum *Nucleocytoviricota* is a diverse viral group that includes families such as *Mimiviridae* and *Phycodnaviridae* (3; 4). *Phycodnaviridae* are mostly lytic viruses characterized by large icosahedral capsids (up to 200 nm in diameter) surrounded by a lipid membrane and contain large dsDNA genomes (up to 600 kb) (3). They play crucial roles in the ocean by regulating the densities of their photosynthetic hosts (phytoplankton) and modulating ecosystem processes (5). Through cell lysis, *Phycodnaviridae* alter the fate of organic carbon and can significantly increase or decrease carbon export to the deep ocean (6; 7; 8; 9). The infection of phytoplankton by phycodnaviruses begins with the attachment of the viral particle to the host cell, followed by the injection and replication of viral DNA and the assembly of viral particles using the host’s cellular machinery. The cycle ends with the lysis of the host cell and the release of new viral particles into the environment (10). Each step of the lytic cycle can be characterized by viral life history traits (LHTs) that modulate the infection cycle, the host-virus population dynamics and the impact of viruses on the ecosystem (11; 12). Given the variability of the ocean environment, the dynamics of phycodnaviruses and phytoplankton are strongly driven by environmental factors that modify LHTs(13). Among these factors, temperature stands out as one of the most critical determinants (14).

Temperature plays a crucial role in shaping virus-host interactions and viral distribution in the ocean by influencing viral LHTs. Key viral LHTs, such as adsorption, latent period, burst size and decay rates, are highly sensitive to temperature changes. Elevated temperatures tend to accelerate viral replication but also increase viral decay rates (15; 16; 17; 14; 18). As a result, temperature-driven changes in these traits impacts virus-host population dynamics. Since virus and host metabolism are interconnected, the viral response to temperature is influenced by how the host reacts to temperature changes (17; 14; 18). A clear example of this can be seen in prasinoviruses infecting the genus *Micromonas* (14; 18): At temperatures below the host’s optimal growth temperature, the viral cycle is optimal, leading to efficient lysis of the host population. However, at lower temperatures, the virus takes longer to complete its cycle, resulting in delayed host lysis. When temperatures get closer or exceed the host’s optimum temperature, the viral lytic cycle is altered, and host cell lysis is significantly reduced (14). It has been suggested that loss of viral infectivity at higher temperatures plays a key role in reducing viral lysis and altering virus-host dynamics (18). On a broader, community-wide scale, viral distribution in marine environments is influenced by temperature gradients, with distinct viral communities thriving in different thermal niches (19; 20; 21).

Understanding how temperature influences viral LHTs and linking these effects to the distribution of the viral community is crucial in the context of global warming. Ocean warming may increase viral production (22; 14; 18; 23; 24), but it can also lead to greater viral decay and loss of infectivity (14; 18). Therefore, understanding the impact of temperature on viral community distribtuion is challenging due to these opposing effects, highlighting the need for interdisciplinary science. In a previous study, we proposed temperature-driven virus distribution patterns based on environmental niches where temperatures are favorable for viral growth (18). Our interdisciplinary approach integrated experimental data with a temperature-driven model and allows us to predict the distribution of viruses and the impact of climate change on the distribution of the virus community. We estimated that the virus distribution will shift towards the poles due to the higher temperatures in the future tropics (18). This study was possible due to experimental studies on the prasinovirus-*Micromonas* response to temperature. This virus-host pair represents an opportunity to develop knowledge on the impact of temperature on marine viruses *in situ* due to their cosmopolitan distribution and the variety of host thermotypes (25; 26).

In this paper, we focus on prasinoviruses that infect members of the dominant green picoalga *Micromonas* and integrate global metagenomes of *Tara* Oceans and Arctic expeditions with a mathematical model to better understand the temperature-driven distribution pattern of marine viruses. We demonstrate that temperature is the primary environmental factor influencing MicV distribution, similar to its host (25). Phylogenetic analysis and viral distribution show that MicV-s are grouped into cryo- and cryo-mesophile communities with distinct distribution patterns. Using our temperature-driven mathematical model, we estimate the distributions of MicV groups and suggest that lysis and infection abilities are good indicators of the presence and absence of lytic viruses in the ocean. Our study emphasizes the value of combining mathematical modeling with experimental and environmental data to enhance our understanding of virus ecology in the ocean.

## 2 Methods

### Phylogeny and Biogeography Analyses

#### Sample Collection

*In situ* samples were collected during the *Tara* Oceans expeditions (26). A non-redundant gene catalog (Ocean Microbial Reference Gene Catalog version 2, OM-RGC.v2) was constructed from 370 metagenomic samples collected during these expeditions (27). An abundance profile of *Nucleocytoviricota*, including MicV, was constructed based on the abundance matrix of the marker gene, family B DNA polymerase (PolB) (28). An abundance matrix of eukaryotes was also generated using metabarcoding data of the V9 region of the 18S rRNA gene (29). From these abundance matrices, samples from the pico-size fraction (0.22–1.6 µm or 0.22–3.0 µm) and the total size fraction (*>*0.8 µm) were used to generate abundance profiles of viruses and eukaryotes, respectively, as these fractions contained higher proportions of targets. Both abundance matrices were normalized to the total number of target communities.

#### Recruitment of MicV Marker Genes

Taxonomic classification of MicV PolB sequences derived from OM-RGC.v2 was conducted using EPA-ng (30), which is based on evolutionary placement algorithms (Supplementary figure S4). To generate the reference tree (backbone) of MicV, we used 22 full-length PolB sequences from MicV isolates and 64 long (*>*700 aa) environmental PolB sequences from OM-RGC.v2. A set of 41 short PolB sequences from MicV isolates with host information (31; 17) was also used to define viral groups in the reference tree to guide taxonomic affiliations. A maximum likelihood tree was constructed using the randomized accelerated maximum likelihood (RAxML) program (version 8.2.12) (32). The environmental MicV PolB sequences were aligned with the reference sequences and then classified into 4 groups (A, B, C, and Pol) based on the maximum likelihood of their placement in the tree. Groups A, B, and Pol were further subdivided into A1, A2, B1, B2, Pol1, and Pol2. A total of 122 sequences were associated with reference MicV-s. We manually checked the taxonomic assignments of these sequences by reconstructing the ML tree and removed 6 of the 122 sequences from downstream analyses, as they were placed far from the reference sequences.

#### Recruitment of *Micromonas* Sequences

The taxonomic assignment of 18S rRNA eukaryote gene sequences was performed using EPA-ng (Supplementary figure S5). The reference ML tree was built using 28 full-length 18S rRNA genes derived from *Micromonas* and an outgroup of Mamiellophyceae (Demory et al., 2018) with RAxML (version 8.2.12). The environmental V9 18S rRNA gene sequences previously assigned to Mamiellophyceae were aligned with the reference sequences and then classified into *Micromonas* groups A (*M. commoda*), B (*M. bravo*), C (*M. pusilla*), and Pol (*M. polaris*). Groups A and B were further subdivided into A1, A2, B1, B2, and B3.

#### Biomes Definition

We defined five biomes according to the latitude (*lat*) and the sample temperature (*T*) in the *Tara* dataset: polar (*lat* ⩾ 60^°^), temperate cold (23^°^ ⩽ *lat <* 60^°^ and *T <* 20°C), temperate warm (23^°^ ⩽ *lat <* 60^°^ and *T* ⩾ 20°C), tropical cold (*lat <* 23^°^ and *T <* 20°C), and tropical warm (*lat <* 23^°^ and *T* ⩾ 20°C).

#### Geographical Viral Community Variation

Community variation among samples was assessed using a Principal Coordinate Analysis (PCoA, function *pcoa* from the package *ape*) based on the Bray-Curtis dissimilarity matrix (function *vegdist* from the package *vegan*). Geographical community variation was verified using a permanova test (function *adonis2* from the package *vegan*) with 9999 permutations between polar and non-polar communities and between non-polar communities only. For each viral phylotype, the optimal temperature was defined as the temperature at which the maximum frequency was observed.,

#### Drivers of Viral Community Distribution

Relationships between environmental variables and viral community distribution were analyzed using a linear regression model (function *lm* from the package *stats*). The environmental variables tested were nominal sample depth (*z*), mixed layer depth (*MLD*), temperature (*T*), salinity (*Sal*), phosphate (*P*), nitrite/nitrate (*N*), silicium (*Si*), and chlorophyll *a* (*chla*). These variables were fitted to the first two dimension scores from the PCoA. The correlation between temperature and the first PCoA dimension was assessed by calculating the Pearson correlation coefficient (function *cor*.*test* from the package *stats*).

#### Phylogenetic Tree Construction

To construct a ML phylogenetic tree of *Micromonas* viruses, we used four sets of PolB sequences: 1) full-length PolB sequences of MicV isolates and their outgroups (Bathycoccus viruses and Ostreococcus viruses), 2) long (>700 aa) environmental PolBs from the OM-RGC.v2, 3) short PolB sequences from MicV isolates with host information (31; 17), and 4) a total of 122 environmental sequences recruited from the OM-RGC.v2 by EPA-ng analysis. These sequences were aligned with MAFFT (ver.7.487) (33) and the aligned sequences were trimmed with trimAl (ver.1.4.1) (34) using default settings. The tree was built with the best-fit substitution model LG+F+G4 and 1,000 iterations of ultrafast bootstrap approximation using IQ-TREE (ver.1.6.12) (35).

#### Network Analyses

We constructed a network based on co-occurrence patterns. We combined the abundance matrices of MicV-s and their hosts. Then, infrequent phylotypes (observed less than 3 samples) and samples (having less than 3 phylotypes) were filtered out from the combined matrix. The resulting abundance matrix was normalized using centered log-ratio (clr) transformation after adding a small pseudocount. An interaction network was inferred using FlashWeave (ver.1.6.2) with sensitive mode and a threshold of *α <* 0.01.

Among 430 significant pairwise associations, 420 positive pairs (97.7%) were used for network visualization using the R package “tidygraph” (https://cran.r-project.org/web/packages/tidygraph).

### Temperature-driven mathematical model

#### Model Description

We used a population model describing interactions between *Micromonas* and its prasinovirus (18). Briefly, susceptible host cells (*S*) can be infected by infectious viral particles (*V*_*i*_) and become infected (*I*). Infection leads to lysis and the release of viral particles into the medium. Virus particles can lose their infectivity and become defective (*V*_*d*_). The dynamics are described as follows:

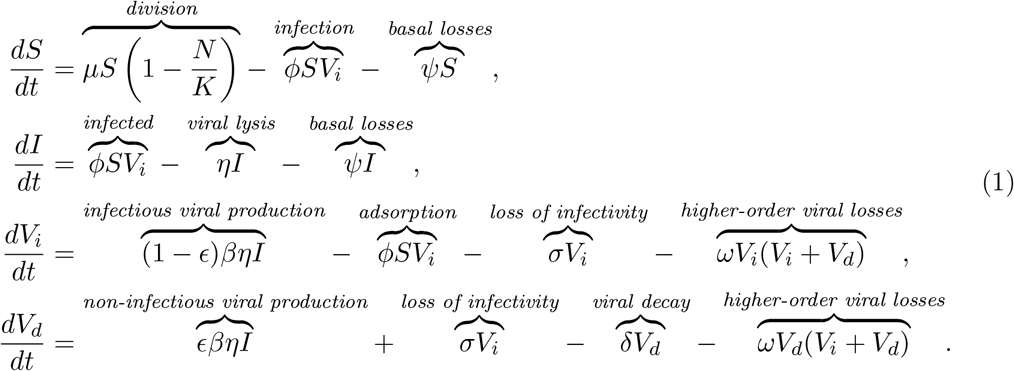

In this model, the interaction between viruses and their host is characterized by life history traits that are drived by temperature. Model parameters are described in the Supplementary Table S2 and in (18).

#### Estimation of Temperature-Driven LHTs

We used parameters estimated by (18) for three virus-host pairs: Mic-A/MicV-A (RCC451/RCC4265), Mic-B/MicV-B (RCC829/RCC4265), and Mic-C/MicV-C (RCC834/RCC4229). Model fits and LHTs as a function of temperature for these three pairs can be found in (18). Additionally, we fitted our model for one virus-host pair characterizing the polar region. We used experimental dynamics of *Micromonas polaris* strain TX-01 infected by virus MpoV45T from (17). Similarly to (18), we used the MATLAB Differential Evolution algorithm (36) to estimate the best parameter set minimizing the error between model fits and experimental data for four temperatures: 0.5, 2.5, 3.5, and 7°C. Host and virus fits are presented in Supplementary Figures S1 and S2, and LHTs as a function of temperature in Supplementary Figure S3.

#### Viral infection and lysis indicators

Following (18), we computed the basic reproduction number associated with viral invasion, the averaged number of newly infections due to the invasion of one virus particle in a virus-free host population, as follows:

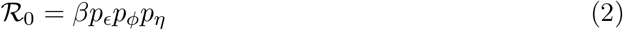

where *β* is the burst size, *p*_*ϵ*_ is the proportion of produced infectious viral particles, *p*_*ϕ*_ is the probability of infection, and *p*_*η*_ is the probability of lysis as follows:

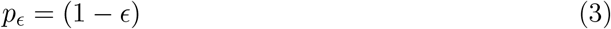

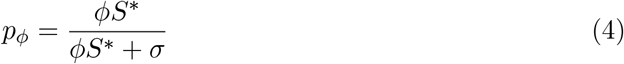

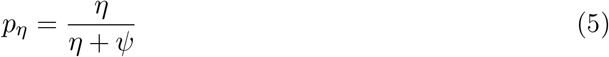

where *ϵ* is the proportion of produced defective particles. *p*_*ϕ*_ is the probability that a virus infects a virus-free host population before losing its infectivity, with *ϕ* being the adsorption rate, *δ* the viral decay rate, and 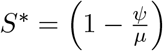 *K* is the disease-free equilibrium calculated by considering no viruses in the environment. Here, *ψ* is the host basal mortality rate, *µ* the host net growth rate, and *K* the host carrying capacity. *p*_*η*_ is the probability that an infected cell lyses before dying due to other mortality processes, with *η* being the lysis rate. More information on parameters can be found in Supplementary Tables S2 and S3.

#### Receiver Operating Characteristic Analysis

To estimate our model’s ability to predict the presence and absence of viruses in the *Tara* dataset, we performed a binary classification analysis using the R package *pROC*. First, for each viral group *i* (A, B, C, and Pol), we classified the data as follows:

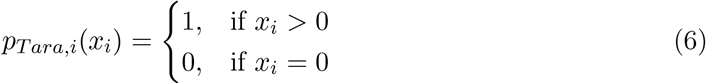

where *x*_*i*_ is the relative frequency of viral group *i*. Second, we computed the probability of lysis and infection, *p*_*η*_ and *p*_*ϕ*_, given the temperature at each *Tara* station. We then computed the Receiver Operating Characteristic (ROC) curves using the function *roc*. We assessed the model’s ability to estimate presence and absence by computing the Sensitivity (or Recall) and Specificity rates as follows:

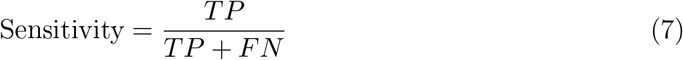

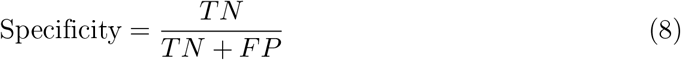

where *TP* is the true positive rate, *TN* is the true negative rate, *FP* is the false positive rate, and *FN* is the false negative rate. We also computed the area under the curve (AUC) and other statistics for our classification as follows:

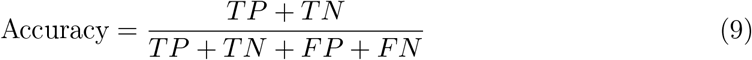

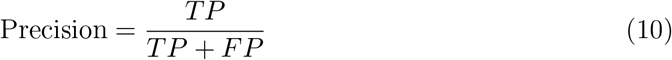

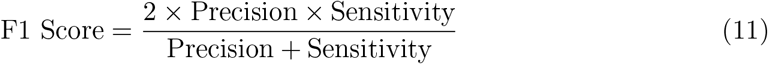

We selected the best thresholds that maximize Sensitivity and Specificity based on the ROC curves for each viral group.

#### Global Distribution Model Estimation

Following (18), we used global monthly averaged SST projections from the NOAA GFDL CM2.1 model (GFDL Data Portal: http://nomads.gfdl.noaa.gov/) run for the years 2010 to 2020. We defined the critical temperature of presence according to the probability of lysis threshold calculated from the ROC analysis for each MicV group. This temperature defined the maximum temperature of presence given the distribution of MicV groups in the *Tara* dataset. Additionally, we calculated the maximum temperature of growth where ℛ_0_ = 1. We mapped the model predictions of presence and absence using the MATLAB package *M*_*Map*_ (37). Model classification abilities were estimated using the ROC results and the classifications of true positive (TP), true negative (TN), false positive (FP), and false negative (FN) samples.

#### Statistical software

Computational analyses were performed using R (version 4.2.3, https://www.r-project.org/, (38)) with Jupyter notebook (https://jupyter.org/, (39)) and MATLAB (version R2023a, https://fr.mathworks.com, (40)).

## 3 Results

### Temperature is the strongest abiotic descriptor of the MicV distribution in the global ocean

We explored the distribution of the MicV community in surface oceans using metagenomic data from the *Tara* Oceans and the *Tara* Polar Circle expeditions (Figure 1 a). After phylogenetic placement and manual curation, we defined 116 PolB sequences as *Micromonas* viruses. MicV sequences were present globally and represented a large proportion of the *Nucleocytoviricota* community in temperate cold and polar biomes (Figure 1b). MicV dominated the *Nucleocytoviricota* community in the Arctic Ocean with a proportion superior to 50% at many stations. In contrast, in warmer regions, MicV sequences represented only a small fraction of the *Nucleocytoviricota* community with proportions less than 5%. Multivariate analysis revealed that MicV communities exhibit dissimilarities between biomes, especially between polar and other regions (Figure 1c). Temperature, salinity, and chlorophyll *a* concentration were the main descriptors of the composition of the MicV community (Figure 1 d). However, temperature was by far the strongest descriptor, explaining 25% of the variance in the MicV composition (Figure 1e and Supplementary Table S1). The MicV community displayed a gradient as a function of temperature, distinguishing between the 5 biomes: Polar, Temperate cold, Temperate warm, Tropcial cold and Tropical warm.

**Figure 1.**
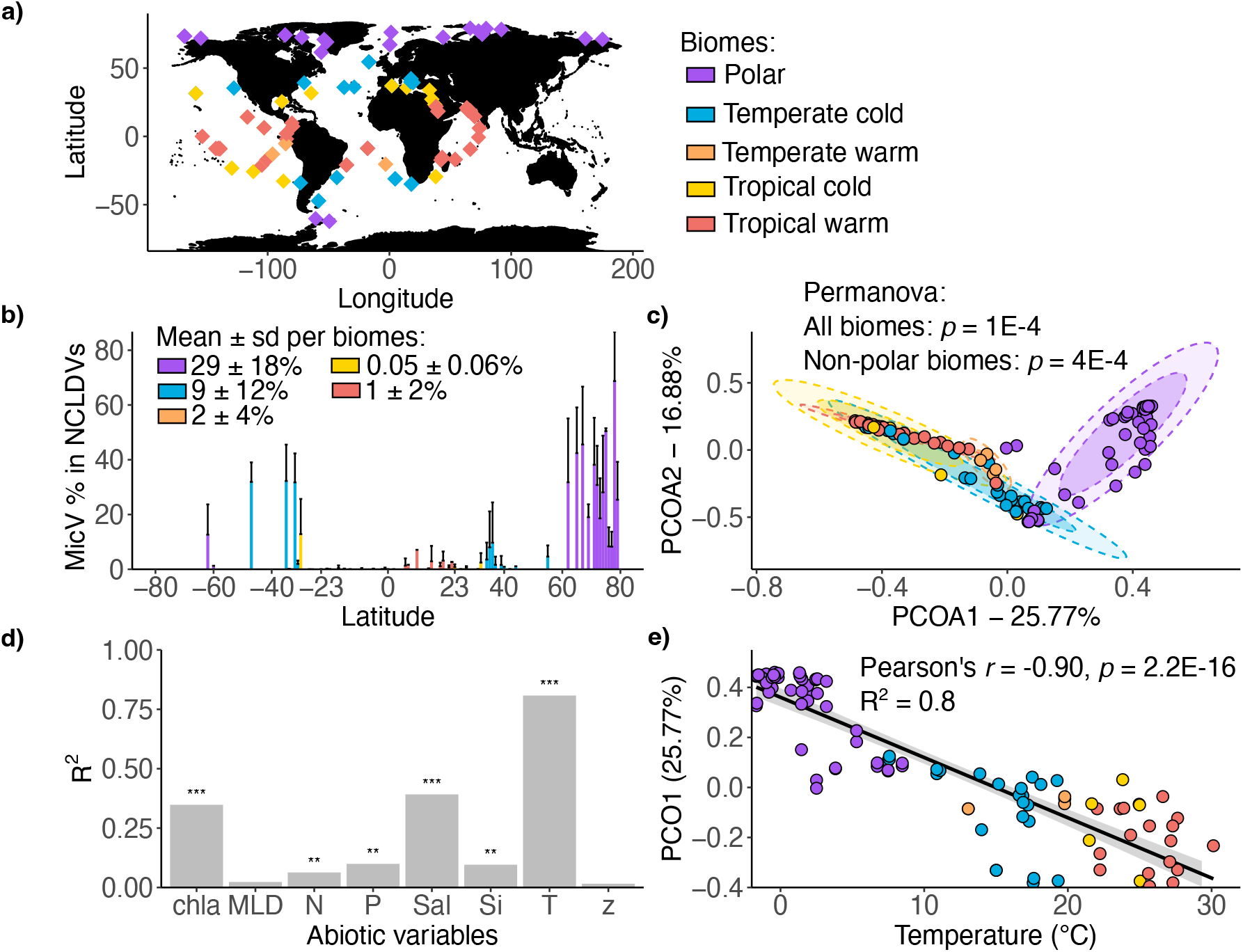
Environmental descriptors of MicV community distribution in global surface ocean. **a)** Location map of *Tara* Oceans expedition samples. **b)** Relative frequencies of MicV within the *Nucleocytoviricota* communities as function of latitude of the sample sites. **c)** Principal coordinate analysis (PCoA) with bray-curtis dissimilarity of the MicV communities. Ellipses represent the 75% and 95% CI of the centroids respectively. **d)** Goodness of fit (*R*^2^) of linear model of the first coordinate of PCoA vs. environmental variables: Chlorophyll a (chla), Mixed Layer Depth (MLD), Nitrates/Nitrites (N), Phosphates (P), Salinity (S), Silicates (Si), Temperature (T) and depth (z). **e)** Linear model of the first coordinate of PCoA vs. temperature.

### MicV groups show either cryophile or cryo-mesophile preferences

We reconstructed a phylogenetic tree of MicV-s using reference and environmental PolB sequences to find relationships between MicV lineages and their thermal environment (Figure 2a). Reference MicV sequences with known host species (31) were used to classify various MicV lineages (Supplementary Figure S4). Viruses infecting *Micromonas pusilla* were found to form a monophyletic group C. In contrary, MicV-s that infect other *Micromonas* species did not form monophyletic groups. Viruses of *Micromonas commoda* were placed in two groups (A1 and A2), those of *Micromonas bravo* were in two groups (B1 and B2), and those of *Micromonas polaris* were in two groups (Pol1 and Pol2). Group A largely dominated the MicV composition with relative frequency within the MicV community over 50% in all temperatures while other groups were present only at temperatures below 20°C (Figure 2b). Group A2 dominated the composition up to 26°C, and group A1 was the only group present at temperatures beyond 28°C. Groups B and Pol represented together between 25 and 40% of the MicV community at low temperatures, and group C was the least frequent MicV group. MicV groups displayed significant variability in their optimal environmental temperature (Figure 2 c). Group A1 had an optimal temperature at 17.07 ± 6.28 (n = 13) with a lower variance compared to A2 (12.91 ± 10.01, n = 55). Group C also had an optimal warm temperature at 13.42 ± 0.52 (n = 5) with a narrow thermal range. Groups B1 and B2 had low optimal temperatures with 4.79 ± 4.77 (n = 18) and 4.97 ± 5.65 (n = 5), respectively. Finally, the Pol1 and Pol2 groups also had low optimal temperatures, but Pol1 had a wider distribution with 7.66 ± 8.65 (n = 15) compared to Pol2 3.92 ± 3.54 (n 5). MicV groups A had a cryo-mesophile profile and were present in lower chlorophyll *a* environments, while groups B and Pol were cryophiles and present in environments with high and low chlorophyll *a*. Group C had a more mesophilic profile, but was present in only a few stations (Figure 2c).

**Figure 2.**
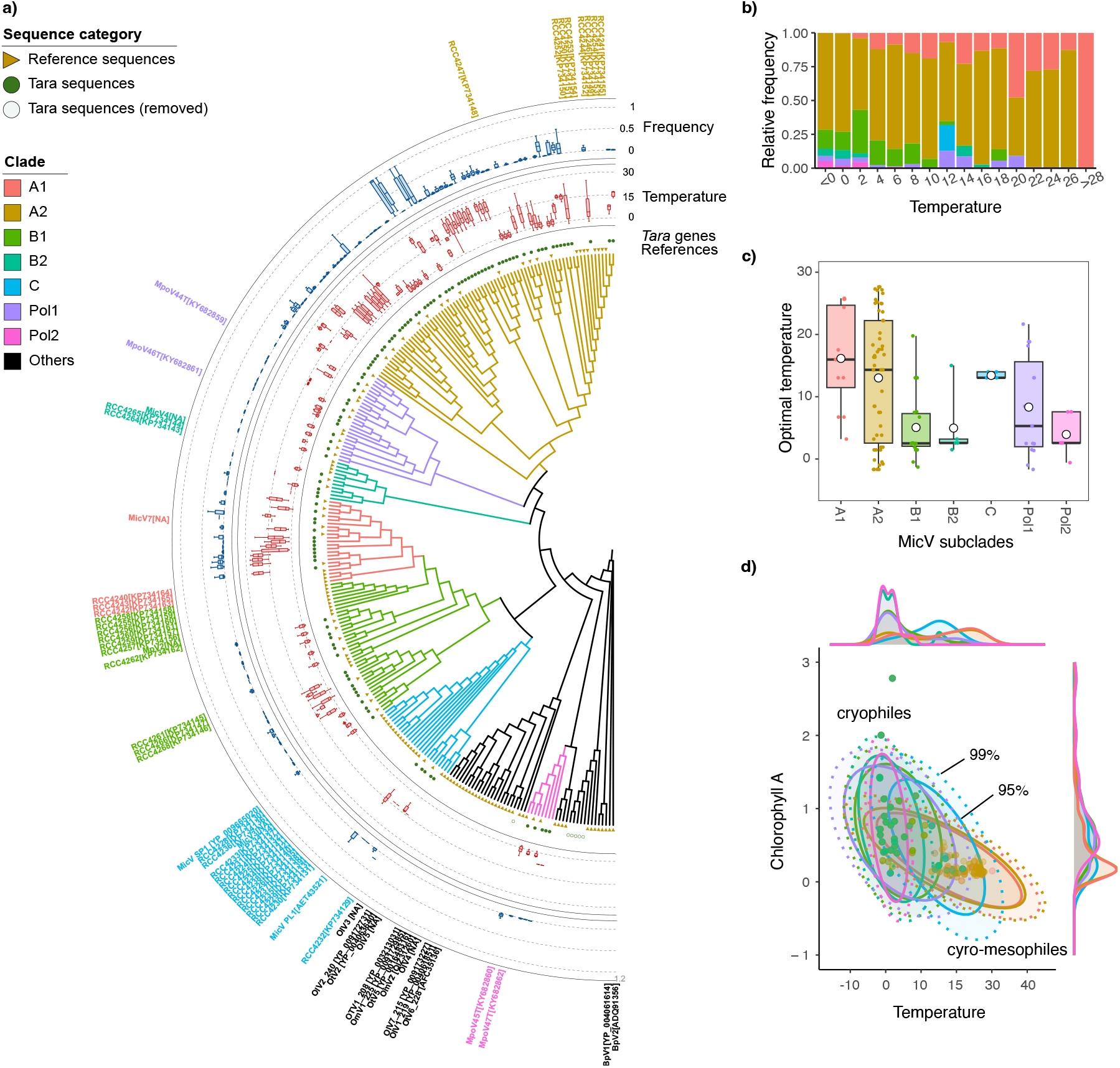
Relation between thermal distribution and MicV phylogeny. **a)** Phylogenetic affiliations of environmental MicV-s. Reference and environmental PolB sequences were shown in triangle and circle marks, respectively, at the first layer. Environmental sequences placed on other virus lineages were removed from the analysis and shown in open circles in this plot. Box plots in the second and third layers represent the ranges of temperature and relative frequency, respectively. The outside layer indicates the phylogenetic positions of MicV strains having clade information. **b)** Relative frequency of MicV subclades as function of temperature. **c)** Optimal temperature of MicV subclades distributions. Boxplot edges are the 25% and 75% quantiles, horizontal black lines are the means, white circle are the medians, and the vertical black lines are the 5% and 95% quantiles. **d)** Representation of the differences between MicV subclade distribution in the space chlorophyll a and Temperature at the isolation site. Ellipses represent the 95% and 99% CI of the centroids for each subclades.

### MicV and their *Micromonas* hosts have similar environmental optimal temperatures

To further assess the relationship between temperature and the distribution patterns of viruses and hosts at the phylotype level, we conducted a co-occurrence network analysis (see Methods for details). Following convention in biogeographic studies of giant viruses and their eukaryotic hosts (28), and in contrast to the interpretation of associations in time series dynamics (41), we interpret positive associations as a potential indicator of interactions (42). Hence, we constructed an interaction network with 101 viruses and 140 hosts. A total of 430 pairs (1.5% of possible pairs) of phylotypes were found to have significant associations, with 420 pairs (97.7%) being positive associations (Figure 3a), including 103 virus-host interactions, 127 virus-virus interactions, and 190 host-host interactions. For all network edges (that is, positive associations), significant positive correlations were found between the optimal temperatures of the connecting phylotypes (Figure 3b). Host-host associations were concentrated in low-temperature regions, while virus-host and virus-virus pairs were more uniformly distributed throughout the temperature range.

**Figure 3.**
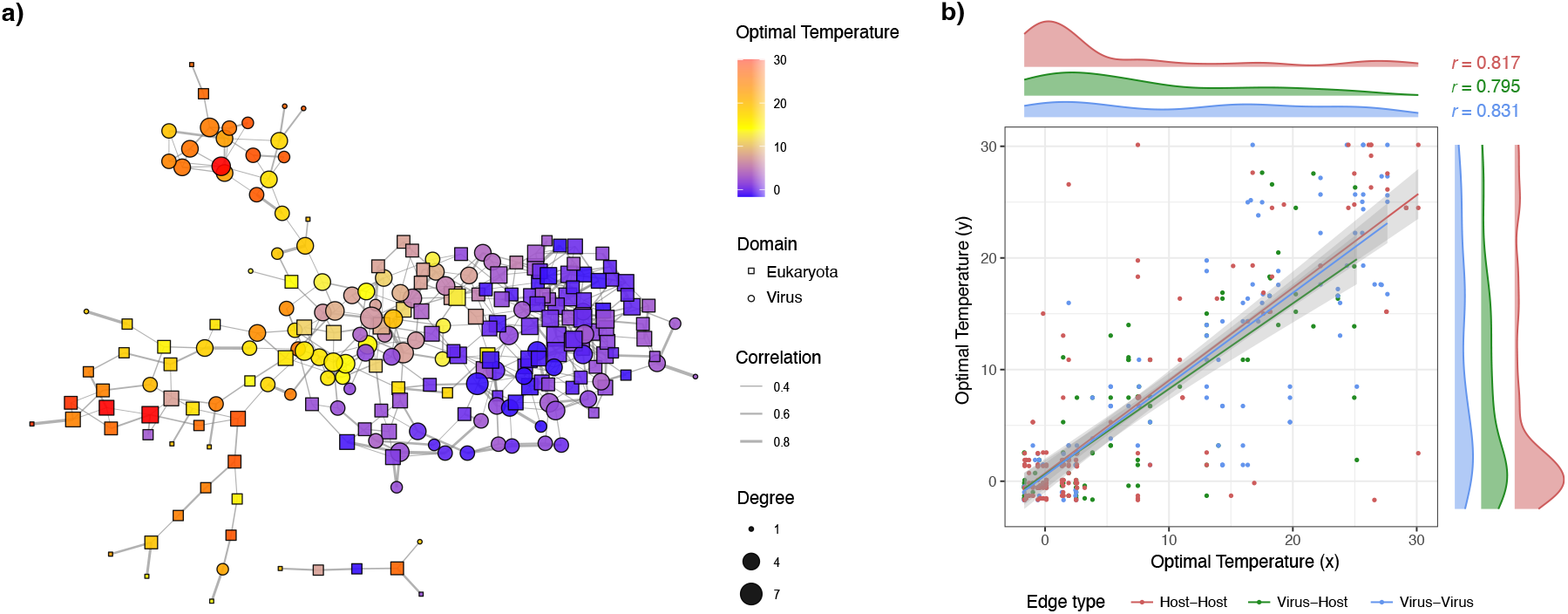
Inference of a virus-host interaction network. **a)** Network is color-coded by the optimal temperatures of viruses (circle) and hosts (square). Edge width and node size indicate correlation coefficient and degree (i.e., the number of connections), respectively. **b)** Relationship between the optimal temperatures of host-host (*n* = 190), virus-host (*n* = 103), and virus-virus (*n* = 127) pairs having positive associations in the network. All the edge types showed significant positive correlation at *p <* 0.01.

### MicV presence and absence are accurately predicted by a temperature-driven model

A mathematical model of temperature impacts on virus-phytoplankton dynamics (18) was used to compare theoretical predictions with the environmental presence of MicV groups in the *Tara* dataset. We extended the model to include the interaction between the MicV group Pol and their host *Micromonas polaris* and integrated experimental data to estimate model parameters (Supplementary Figures S1 and S2). We obtained 4 parameterizations characterizing the interactions of the 4 MicV groups (A, B, C, and Pol) with their host. We used the probability of lysis, the probability of infection and the proportion of infectious produced virions as indicators of the presence and absence of MicV groups and estimated the threshold values by performing Receiver Operating Characteristic (ROC) analyses (Figure 4). The probability of lysis and infection and the proportion of infectious virions were good indicators of the presence and absence of MicV groups A, B, and Pol, given the ROC statistics with precision superior to 70% (MicV A) and 80% (MicV B and Pol) specificity and sensitivity (left panels in Figure 4). The 3 indicators performed poorly in predicting the distribution of MicV group C with accuracy falling around 50% specificity and sensitivity. It is likely that the model failed to predict the distribution of MicV group C due to the small number of sequences in the *Tara* data (only found in 10 stations of 226), but also because viruses from this group have longer latent periods (up to 30 hours), which could reduce the lysis signal.

**Figure 4.**
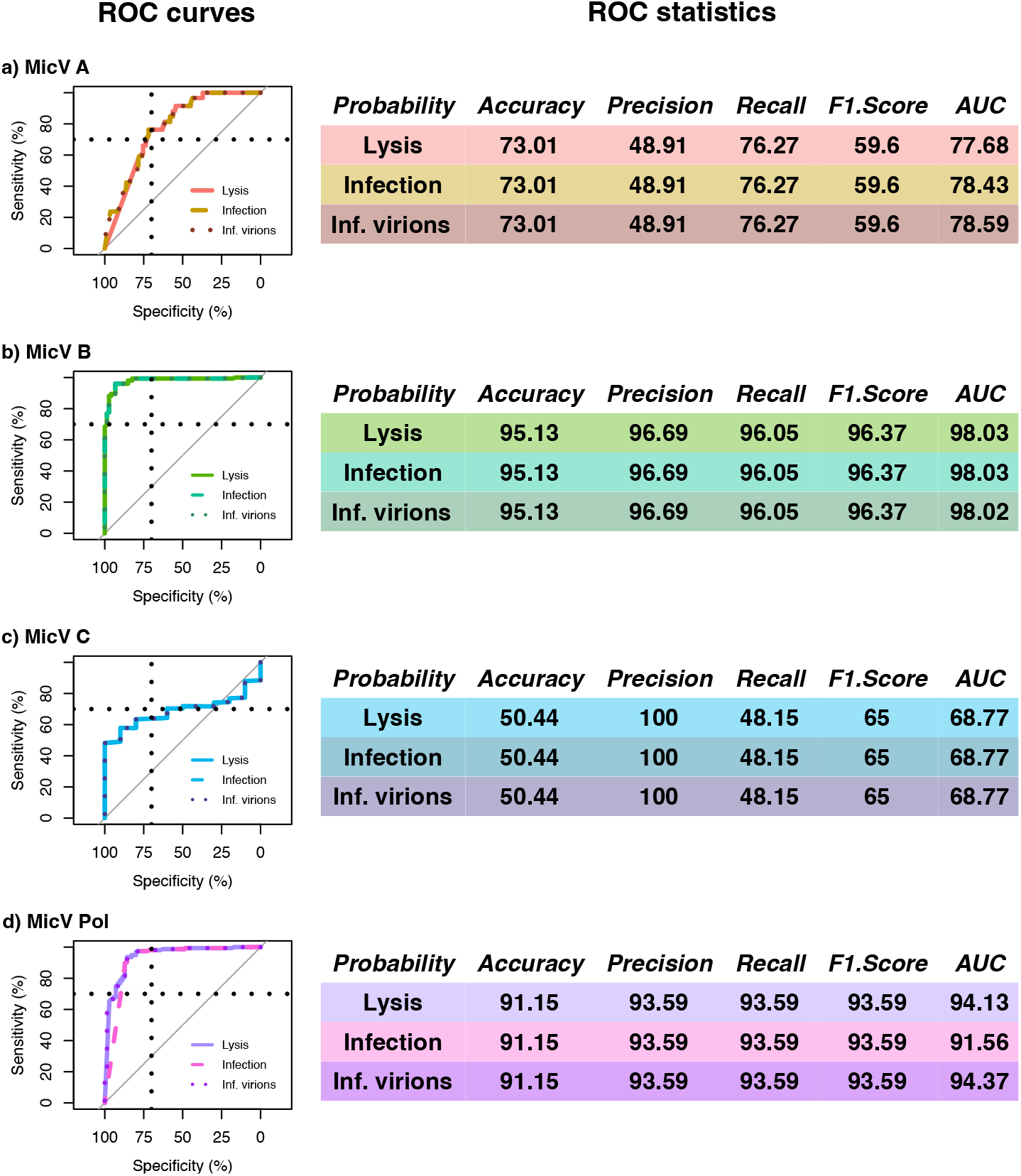
ROC analysis using the probability of lysis and of infection and the proportion of infectious produced virions estimated from mathematical modeling for MicV groups **a)** A, **b)** B, **c)** C and **d)** Pol. Left panels are the ROC curves for probability of lysis (colored solid line), probability of infection (colored medium-dashed line) and proportion of infectious produced virions (colored small-dashed line). Vertical and horizontal dashed black lines are the 70% threshold of Specificity and Sensitivity. Gray 1:1 curve represents models that cannot distinguish between presence and absence in data. Right panels are ROC statistics with Accuracy (how many positive and negative observations are correctly classified), Precision (how many positive estimations are actually positive), Recall or Sensitivity (how many positive observations are classified correctly), F1 score (harmonic mean of precision and recall) and AUC (area under the curve).

### Global model prediction reveals the importance of lysis as a driver of the MicV distribution

Given the similar predictions of our 3 indicators, we used the probability of lysis as a predictor of the presence or absence of MicV in the global surface ocean (Figure 5). We estimated maximum temperature thresholds from the ROC analysis and compared them to the maximum temperature of invasion fitness, when ℛ_0_ *>* 1 for the four MicV groups A, B, C and Pol (Figure 5 left panels). The threshold temperature for group A was similar to the maximum temperature of invasion (23.1°C and 23.5°C, respectively). For MicV group B, we found significant differences between the two temperatures, with 15.1°C and 27.4°C for the threshold and invasion temperatures, respectively. The differences were less significant for group C and Pol, with 21.2°C and 25.6°C for the former and 10.3°C and 13.9°C for the latter. We mapped the presence and absence regions according to the temperature threshold for the four groups and quantified the classification ability of our model by comparing the *Tara* data with our model estimates (Figure 5 middle and right panels). Our model estimated the presence of MicV group A from the polar region to more than 40^°^ latitude with an accuracy of 73% for true positive and true negative samples. Specifically, 53% of the samples were classified as true positive and 19.9% as true negative, while 6.2% and 20.8% were classified as false positive and false negative, respectively. The false negative samples were distributed mainly in the warmer tropical regions. The distribution of group B ranged from the polar region to a maximum of 40^°^ latitude, with 95.1% of the samples having a good classification. True positive samples represented 30.5% and true negative 64.6%, while only 2.7% and 2.2% were classified as false positive and false negative. Similarly to group A, group C had a wide distribution from the polar region to more than 40^°^ latitude, but the precision of our model was poor. True positives represented only 4% of the samples and true negative 47.8%, while 47.8% of the samples were classified as false positive and 0.4% as false negative. False positive samples were distributed mainly in cold environments beyond 40^°^ latitude. The Pol group had an accuracy of 90.7% and was distributed up to 40^°^ latitude. True positive samples represented 26.1% and 64.6% were classified as true negative. False positive and negative accounted for only 4.4% and 4.9%, respectively.

**Figure 5.**
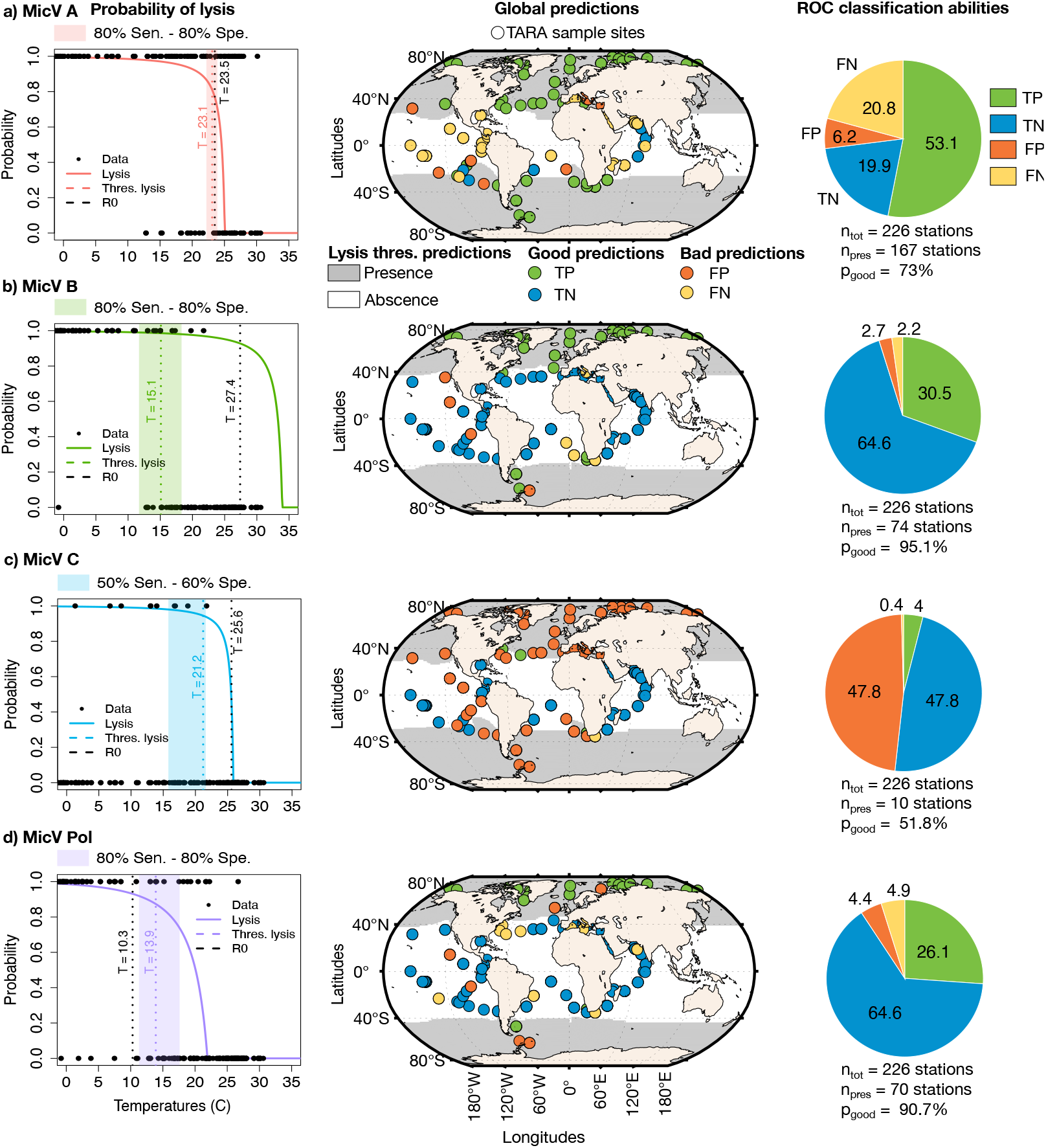
Model estimation of MicV global distribution based on the probability of lysis for MicV groups **a)** A, **b)** B, **c)** C and **d)** Pol. Left panels are the probability of lysis (solid line) as function of temperature. Black dots are *Tara* presence and absence data. We considered presence when the relative frequency of each group is positive and absence when it is null. Vertical dashed black and colored lines are the temperature threshold estimated using R0 and probability of lysis. Shaded areas represent the threshold temperature for the optimum sensitivity (Sen) and specificity (Spe) with 70% Sen - 70% Spe for group A, 80% Sen - 80% Spe for groups B and Pol and 50% Sen - 60% Spe for group C. Middle panels are the global map estimation of presence and absence of MicV groups. Gray areas are the presence regions estimated with the best threshold for the probability of lysis. Dots are *Tara* samples with colors based on their classification: True Positive (TP – green), True Negative (TN – blue), False Positive (FP – orange) and False Negative (FN – yellow). Right panels are the classification percentage for TP (green), TN (blue), FP (orange) and FN (yellow). *n*_*tot*_ is the total number of *Tara* samples, *n*_*pres*_ is the number of *Tara* samples that have presence observations and *p*_*good*_ is the percentage of good estimation of presence and absence (TP+TN) by our model.

## 4 Discussion

In this study, we explored the temperature-driven biogeographic pattern of viruses infecting the picoeukaryote *Micromonas* by integrating environmental metagenomics and mathematical modeling. We showed that temperature is the main driver of the distribution of the MicV community in the global ocean; specifically, we identified cryophile and cryo-mesophile phylogenetic viral groups, following the thermal niches of their hosts. Using a temperature-driven dynamical model of host-virus interactions, we estimated the viral abilities to infect and lyse their host for each viral group as a function of environmental temperatures and predicted the presence and absence of MicV groups. Our modeling predictions matched the presence and absence patterns for most viral groups in the *Tara* dataset.

Our findings are in line with previous studies showing that temperature is one of the main drivers of dsDNA giant virus (28; 21), dsDNA phage (29), jumbo phage (43) and RNA (44) and DNA virome biogeography (19; 20). Interestingly, we found a strong difference between the composition of the polar and non-polar MicV community and that MicV dominates the *Nucleocytoviricota* sequences in the Arctic ocean. Meng and colleagues (21), also reported that the cold temperatures of the Arctic Ocean act as an ecological barrier that separates the polar and non-polar giant virus communities. This ecological barrier might increase viral speciation and represent a hot spot for virus diversity in the Arctic (29; 21). Temperature not only separates polar and non-polar MicV communities but also describes a gradient in their community composition from temperate cold to tropical warm water, explaining up to 23% of the variance in our dataset. Other abiotic factors significantly influenced the composition of MicV, such as chlorophyll *a* and salinity, but had a lower impact. Chlorophyll *a* is often used as a rough proxy for phytoplankton biomass (45; 46) and suggests that host abundances are also driving the biogeography of MicV. Host abundances are highly variable in coastal areas where chlorophyta (especially *Micromonas*) often dominate the phytoplankton community (47; 48), suggesting that they might play an important role in the dynamics of MicV (31).

Despite the strong role of temperature in shaping the distribution of MicV-s, phylogenetic groups cannot be characterized as true thermotypes, as is the case for their host *Micromonas* (25). MicV-s infecting similar hosts were identified in different groups having distinct evolutionary origins in their phylogeny. For example, subgroup A1 is genetically closer to subgroups B1 and C while subgroup A2 is closer to Pol1. This result does not necessarily mean that virus thermotypes do not exist, but that the taxonomic level of our analysis might be insufficient or that the ability of some viral groups to infect different host thermotypes is certainly blurring the signal (31). In addition, viral groups B and Pol can be characterized as cryophilic viruses with an average temperature distribution lower than 10^°^ C, while group A has a wide thermal distribution extending to temperatures higher than 25°C. Within each group, we also identify various thermal optima, suggesting that genetically closely related viruses can have different responses to temperature. Within group A, subgroup A2 has a wide temperature distribution, and dominates the MicV community from 0^°^ to 26°C. Subgroup A1 is not present lower than 10°C, suggesting that it is more adapted to warm environments. The fact that model predictions include high false negative rates for MicV group A distribution prediction in the tropics, might suggest that MicV group A has a higher evolution ability at high temperature and can invade warmer environments. Our findings highlight the complexity of viral thermal responses and suggest that further explorations of the mechanisms driving these patterns are necessary.

Temperature affects viral fitness and distribution through various biological processes, with different impacts at low and high temperatures. At low temperatures, viruses degrade more slowly, suggesting that viral fitness is mostly limited by the host cell’s metabolism and lower growth rate (14; 18). Factors like the variability of *Micromonas* species in growing at low temperatures (25) and abiotic factors such as salinity affect how well the host can support viral replication (49). Specific patterns of infection among virus groups (31) and the impact of coinfection (multiple viruses infecting the same cell) (50) could also explain why certain virus groups, such as MicV group A2, dominate, while others are less prevalent. In contrast, at higher temperatures, rates of viral particle degradation and loss of infectivity are high, suggesting that viral fitness is mostly driven by degradation processes (14; 18). The mechanisms driving viral loss of infectivity response to temperature are not yet well characterized. Temperature can increase the virus polymerase error rate (51), resulting in the production of defective viral particles (18; 52). These defective particles can compete with wild-type virus, potentially reducing their fitness and reducing the ability of viruses to extend their thermal niches in warmer environments. The distribution of MicV-s suggests that these viruses may adapt and evolve more easily to colder rather than warmer environments as previously proposed (18; 21) - though understanding virus evolvability *in situ* remains an open challenge.

Our integrative approach allows for the accurate prediction of the presence of the majority of MicV groups. The probability of lysis and infection and the proportion of infectious produced virions emerge as strong predictors of virus distribution. MicV-s are mostly lytic viruses (53), and therefore, the presence of viral genetic signatures in metagenomes increases with the virus ability to infect and lyse their host. The difference between the temperature threshold (from the ROC analysis, *T*_*realized*_) and the maximum temperature of viral growth (when ℛ_0_ = 1, *T*_*fundamental*_) suggests a discrepancy between the realized and fundamental thermal niches, driven by other processes such as biotic interactions (54). Notably, MicV group A’s *T*_*realized*_ approaches the maximum temperature limit of *T*_*fundamental*_, while MicV group B has a *T*_*realized*_ that is 12°C lower than its *T*_*fundamental*_. Both MicV groups A and B can infect *Micromonas commoda* and *bravo*, resulting in competition for susceptible hosts (31). This result suggests that MicV group A might have higher competitive fitness than MicV group B and can extend its realized niche to the maximum theoretical limits of its fundamental niche. In contrast, MicV group B is limited to a lower maximum temperature, where the competitive pressure from MicV group A (especially subgroup A1, which is not present at low temperatures) may be reduced. MicV group A might have better competitive fitness at high temperatures by reducing viral loss of infectivity, minimizing the production of defective particles through cell lysis (18), or better managing co-infection. Indeed, a recent study proposed that interspecific co-infection drives virus-virus competition in *Micromonas* viruses and that shifts in temperature alter competition outcomes (50). Our study highlights the importance of investigating the impact of temperature on interspecific competition mechanisms.

Despite its simplicity, the model is able to predict the presence and abscence of viruses across the global oceans, albeit with caveats. First, on the experimental side, our model is parameterized on only four virus-host pairs, and we lack experimental studies investigating the intraspecific temperature response within each thermal group. Therefore, we may be missing potential intraspecific variability that could be reflected by some false positive and negative rates in our predictions. To extend our approach to other phytoplankton and virus genera, we also need experimental studies investigating the impact of temperature on virus-host dynamics at different taxonomic levels. Secondly, on the observational side, the *Tara* dataset is biased toward the Northern Hemisphere, especially for the polar region, while the Southern Hemisphere is underrepresented (27). Increasing sampling in the Antarctic region will help to achieve better estimations and validations of virus biogeography. Furthermore, the low presence of MicV group C suggests that our reference sequences may not be large enough to recruit target species from the metagenomes. This highlights the need to expand the full length of PolB sequences, other marker genes, or genomes having host information for better distinguishing intra-genus and intra-species virus groups.

In conclusion, our study represents an attempt to link virus temperature responses to virus biogeography using an integrative mechanistic approach rather than a solely correlative one (55). We find that temperature is central to shape virus ecology and biogeography, but also emphasize that interdisciplinary studies are needed to gain a better understanding of the impact of other environmental factors and feedbacks on virus ecology.

## Acknowledgments

We thank scientists and crew members of the Tara Oceans and Tara Oceans Polar Circle cruise, as well as the Tara Expeditions Foundation.We also thank Dr. Douwe Maat and Prof. Corina Brussaard for providing their original Arctic MpV experimental data and Marian Dominguez-Mirazo for code review. Finally, we thank Jeremy Seurat and members of the Weitz group at the University of Maryland, the Ogata laboratory at the University of Kyoto and the GENOPHY axis at LBBM for our discussions on host-virus interactions, statistical and modelling analyses.

## Supplemental Information

**Table S1.**
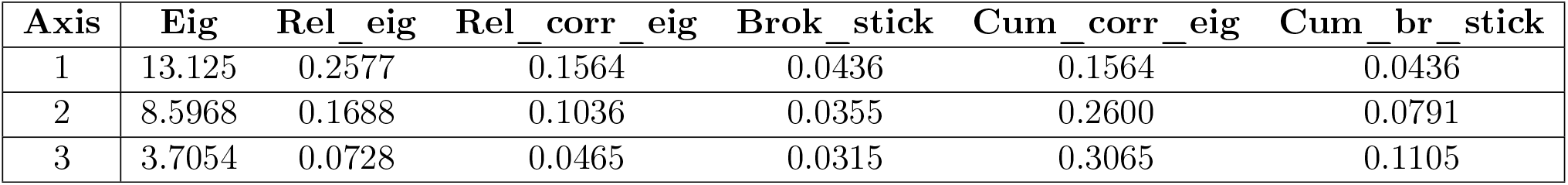
Scores and metrics for the 3 first axes of the PCoA of figure 1c.

**Table S2.**
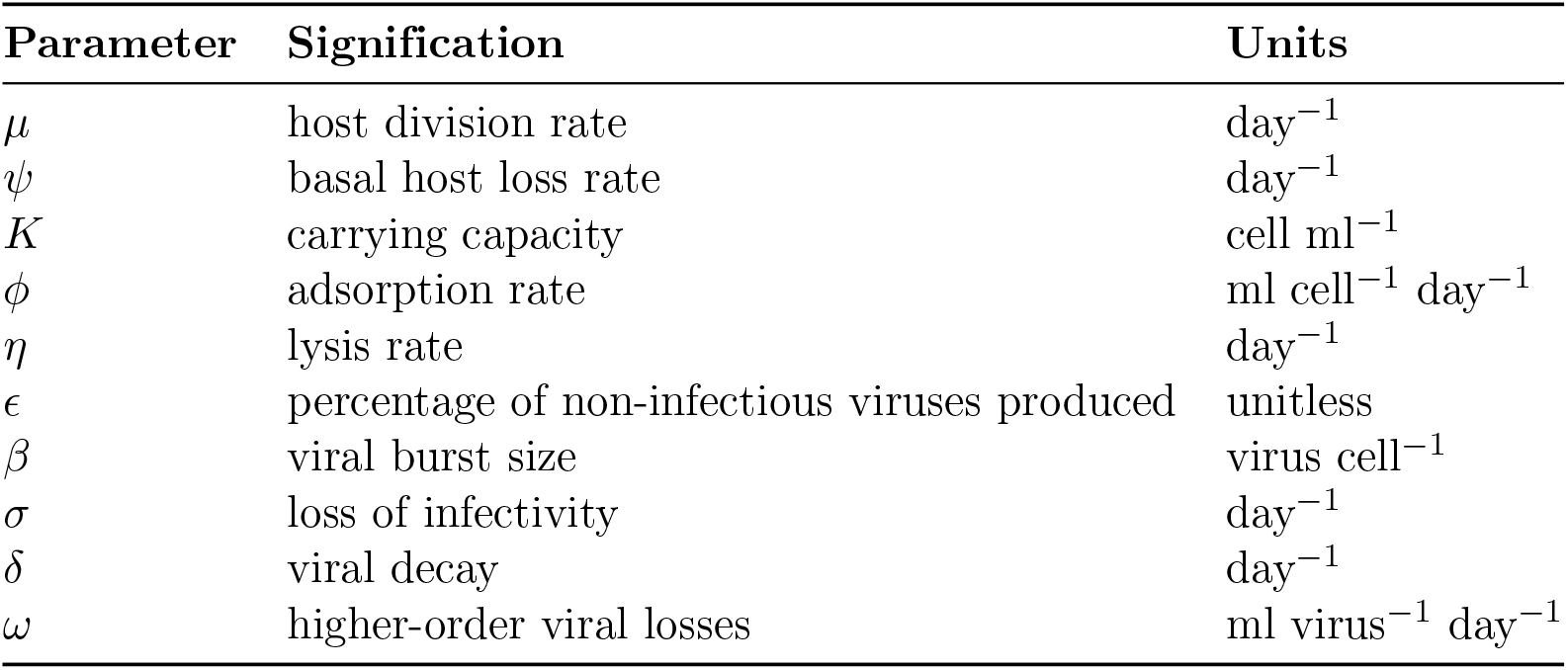
Information about parameters used in the model from (18).

| Parameter | Signification | Units |
| --- | --- | --- |
| $\mu$ | host division rate | day<sup>-1</sup> |
| $\psi$ | basal host loss rate | day<sup>-1</sup> |
| $K$ | carrying capacity | cell ml<sup>-1</sup> |
| $\phi$ | adsorption rate | ml cell<sup>-1</sup> day<sup>-1</sup> |
| $\eta$ | lysis rate | day<sup>-1</sup> |
| $\epsilon$ | percentage of non-infectious viruses produced | unitless |
| $\beta$ | viral burst size | virus cell<sup>-1</sup> |
| $\sigma$ | loss of infectivity | day<sup>-1</sup> |
| $\delta$ | viral decay | day<sup>-1</sup> |
| $\omega$ | higher-order viral losses | ml virus<sup>-1</sup> day<sup>-1</sup> |

**Table S3.** Hyper-parameter values of the temperature-driven functions for the polar host-virus pair calibrated using data from (17). Hyper-parameter values used for the 3 other pairs can be found in (18).

| Hyper-parameter | Mic-Pol/MicV-Pol | Units |
| --- | --- | --- |
| $A1$ | $1 \cdot 10^{11}$ | day<sup>-1</sup> |
| $E1$ | 7260.7 | °K |
| $A2$ | $1.99 \cdot 10^{24}$ | day<sup>-1</sup> |
| $E2$ | 16296 | °K |
| $K$ | $1 \cdot 10^{10}$ | cell ml<sup>-1</sup> |
| $\phi_K$ | $2.63 \cdot 10^{-7}$ | ml (cell day)<sup>-1</sup> |
| $T_\phi$ | 291.83 | °K |
| $\phi_r$ | 0.1 | °K<sup>-1</sup> |
| $s1$ | $2.09 \cdot 10^9$ | day<sup>-1</sup> |
| $d1$ | 5705.3 | °K |
| $s2$ | $3.05 \cdot 10^{24}$ | day<sup>-1</sup> |
| $d2$ | 16007 | °K |
| $b1$ | 99.534 | day |
| $T_\epsilon$ | 280.5 | °K |
| $\epsilon_r$ | 9.2464 | unitless |
| $\sigma_1$ | $5.02 \cdot 10^{22}$ | day<sup>-1</sup> |
| $\delta_1$ | 0.49447 | day<sup>-1</sup> |
| $\delta_2$ | 0.47675 | °K |
| $\omega$ | $1.49 \cdot 10^{-10}$ | ml (virus day)<sup>-1</sup> |

**Figure S1.**
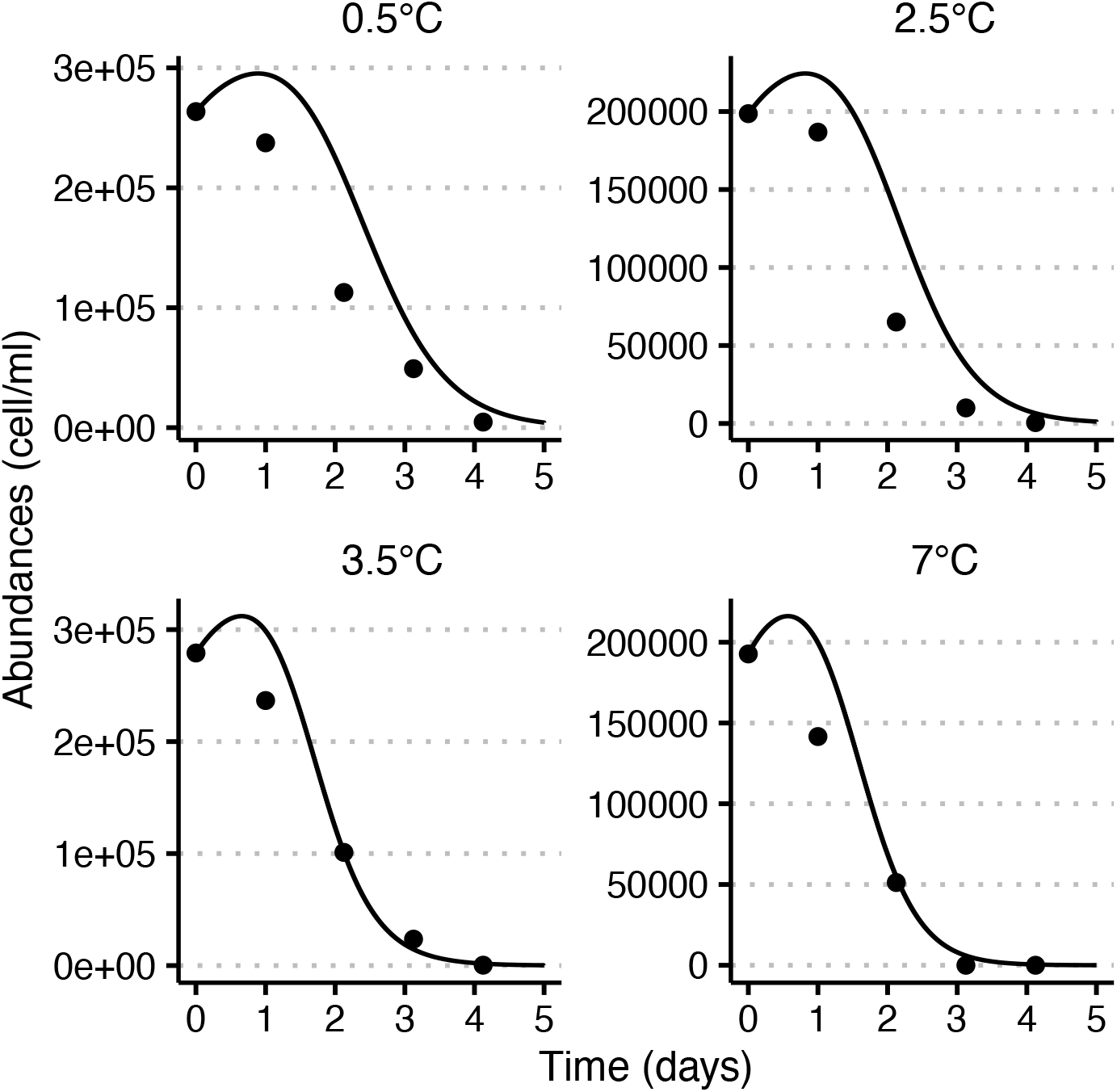
Model-data fitting for the polar virus-host pair. Host dynamics at 0.5, 2.5, 3.5 and 7°C. Solid black lines are the model fits and black dots the data from (17).

**Figure S2.**
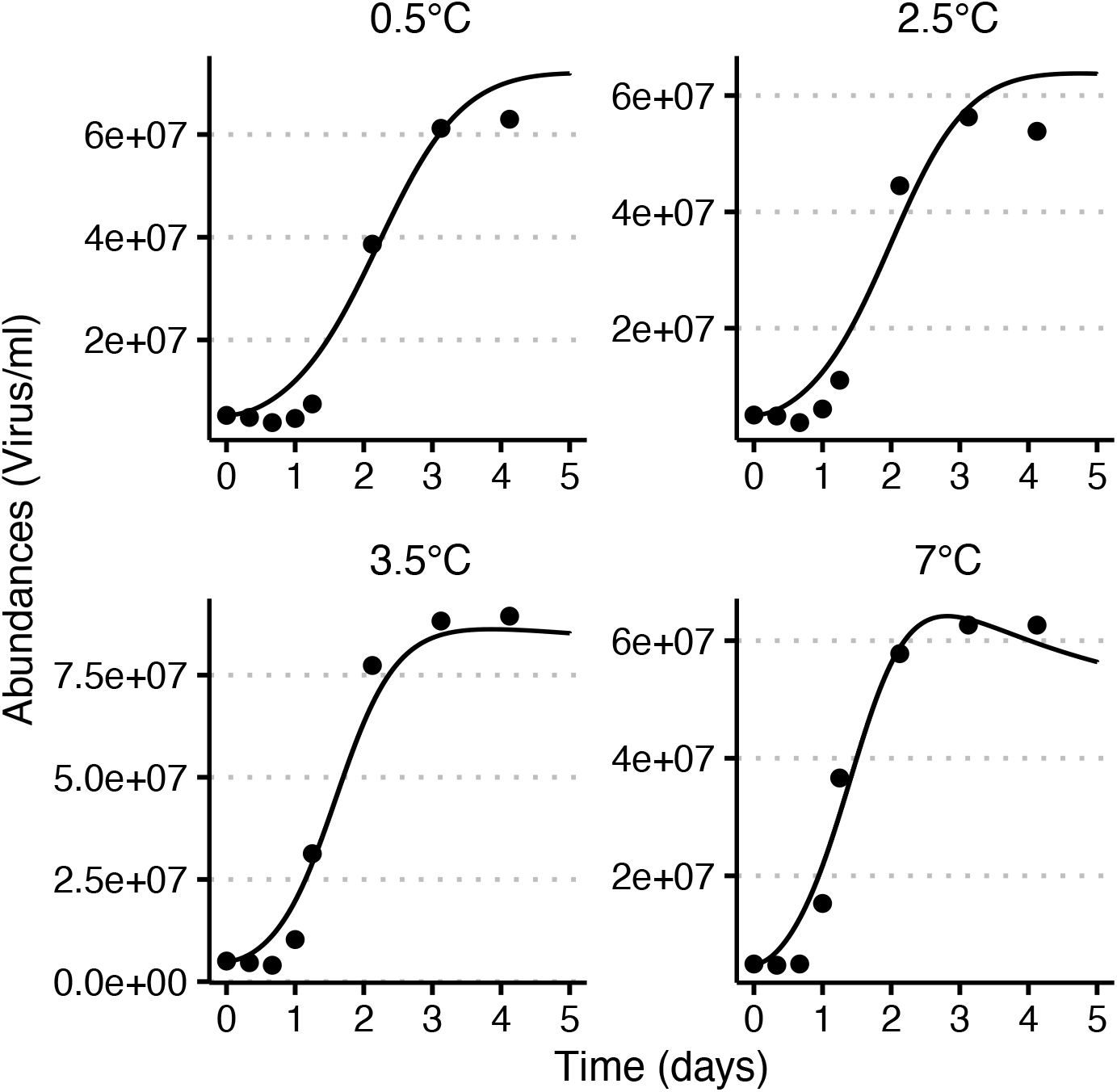
Model-data fitting for the polar virus-host pair. Virus dynamics at 0.5, 2.5, 3.5 and 7°C. Solid black lines are the model fits and black dots the data from (17).

**Figure S3.**
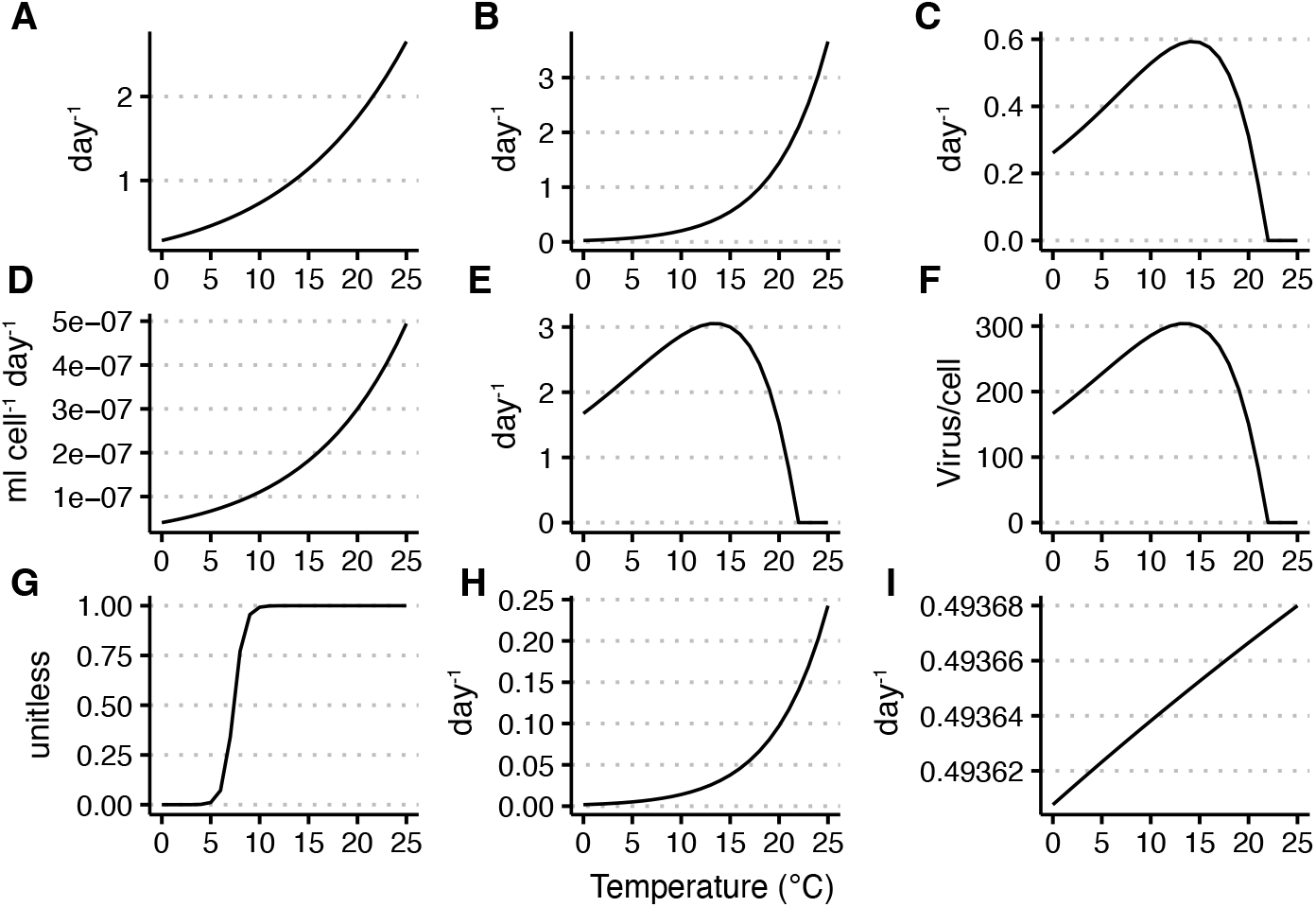
Temperature-driven hyperfunction for the host and virus life history traits estimated from the model-data fitting. A) Host gross growth rate, B) Host mortality rate, C) Host net growth rate (gross growth rate minus mortality), D) Viral adsorption rate, E) Viral lysis rate, F) Viral burst size, G) Proportion of produced non-infectious viral particle, H) Viral particle loss of infectivity and I) Viral degradation rate. More details on the hyperfunction equations can be found in (18) and in the github repository.

**Figure S4.**
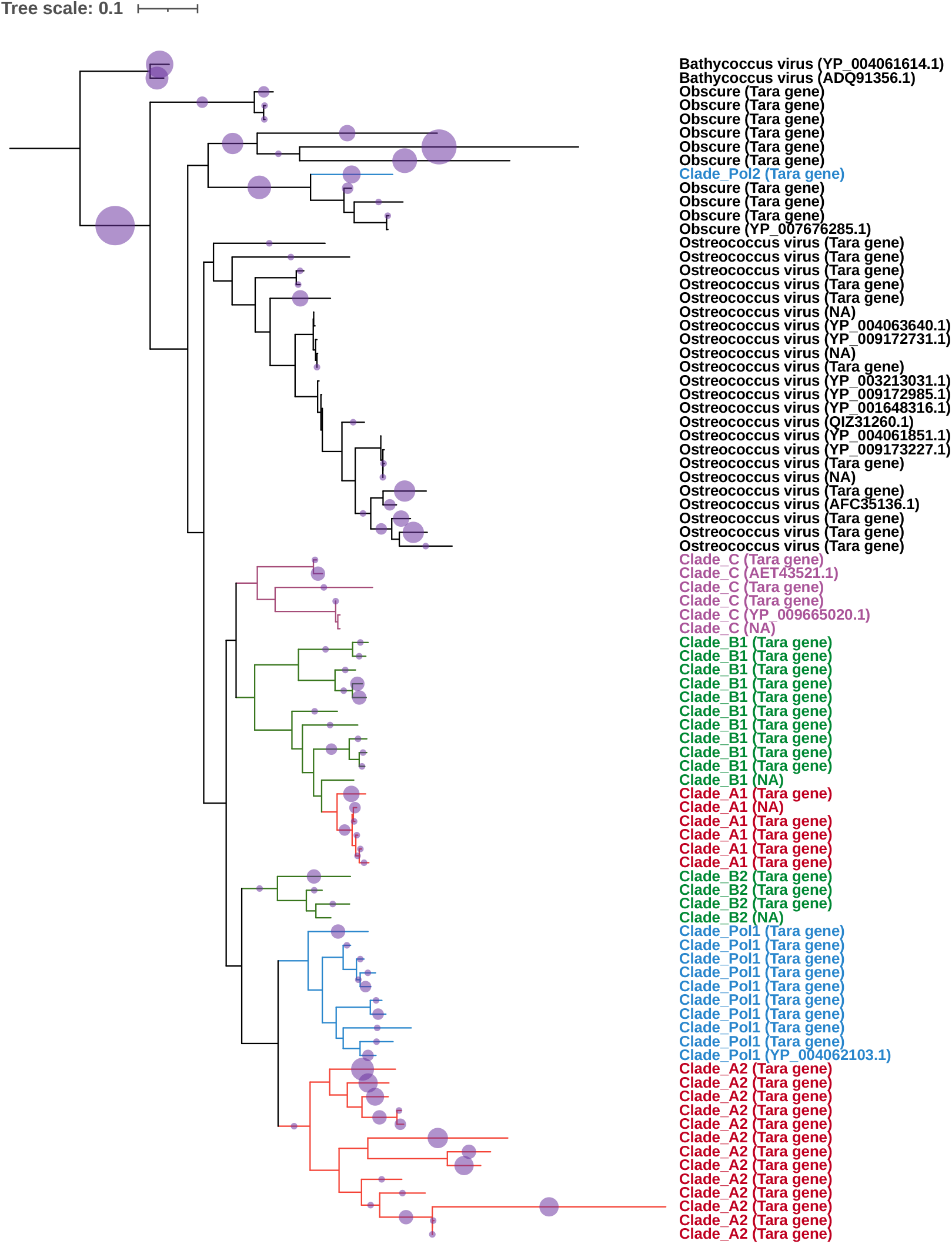
MpV Pylogenic tree of full-length or long (>700 aa) PolB sequences of Prasinoviruses used as the EPA-ng analysis. Taxonomic affiliations were confirmed by the previously published short sequences in (31; 17).

**Figure S5.**
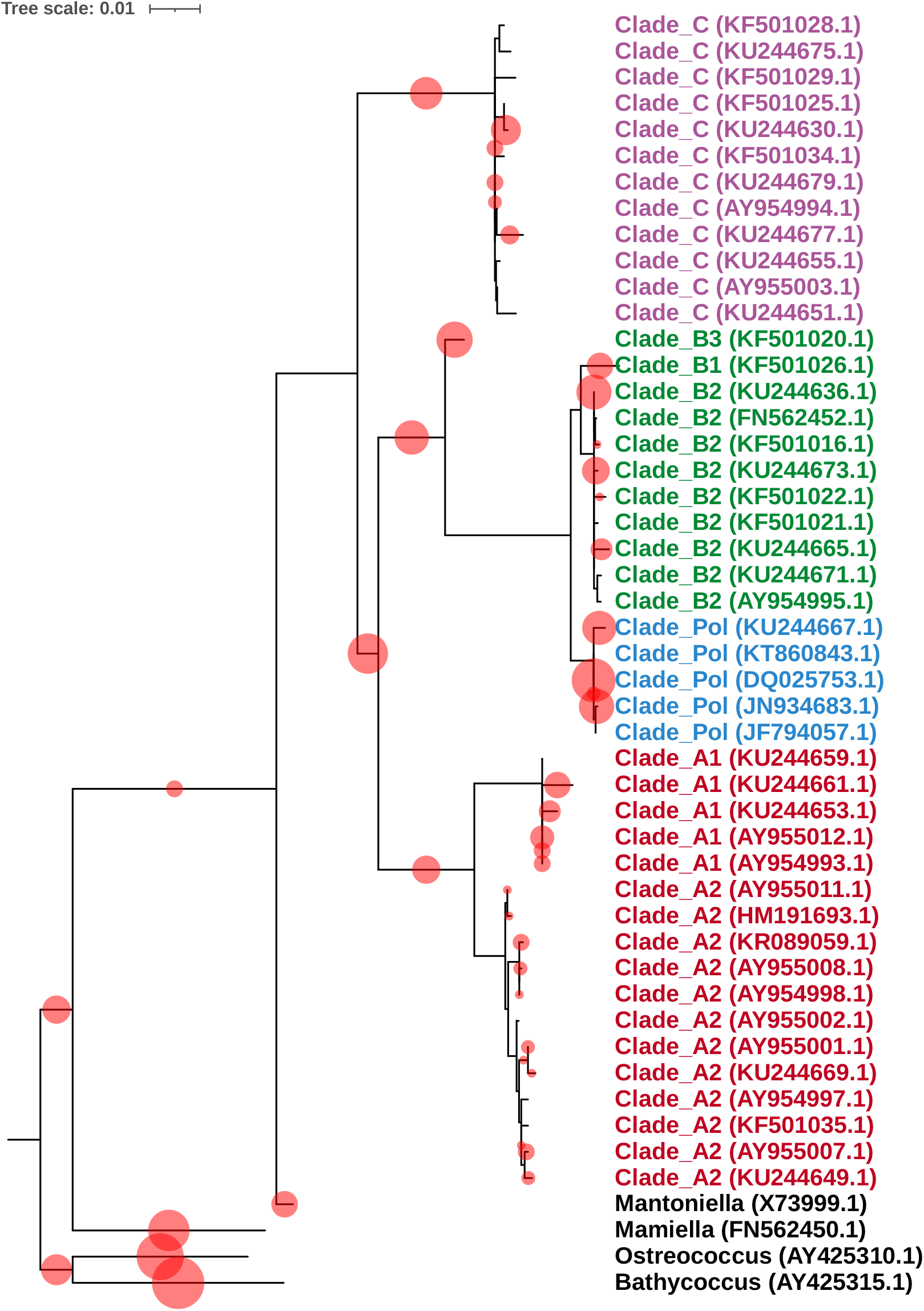
Micromonas Pylogenic tree of full length 18S rRNA gene sequences of Micromonas used as the EPA-ng analysis. Taxonomic affiliations were confirmed by the previously published short sequences in (25).

